# WDR47 facilitates ciliogenesis by modulating intraflagellar transport

**DOI:** 10.1101/2022.04.10.487758

**Authors:** Chun-Xue Song, Xian-Ting Zeng, Wan-Xin Zeng, Xia-Jing Tong, Qian Li

## Abstract

Cilia are conserved organelles found in many cell types in eukaryotes, and their dysfunction causes defects in environmental sensing and signaling transduction; such defects are termed ciliopathies. Distinct cilia have cell-specific morphologies and exert distinct functions. However, the underlying mechanisms of cell-specific ciliogenesis and regulation are unclear. Here we identified a WD40-repeat (WDR) protein, WDR47/NMTN-1, and show that it is specifically required for ciliogenesis of AWB chemosensory neurons in *C. elegans*. WDR47/NMTN-1 is expressed in the AWB chemosensory neuron pair, and is localized at the basal body (BB) of the AWB cilia. Knockout of *wdr47/nmtn-1* causes abnormal AWB neuron cilia morphology, structural integrity, and induces aberrant AWB-mediated aversive behaviors. We further demonstrate that *wdr47/nmtn-1* deletion affects movement of intraflagellar transport (IFT) particles and their cargo delivery in AWB neurons. Our results indicate that WDR47/NMTN-1 is essential for AWB neuron ciliary morphology and function, which reveal a novel mechanism for cell-specific ciliogenesis. Since WDR47/NMTN-1 is conserved in mammals, our findings may help understand the process of cell-specific ciliogenesis and provide insights for treating ciliopathies.

## INTRODUCTION

Cilia are microtubule-based sensory organelles that are found throughout most eukaryotes (Pedersen, Schroder et al. 2012). They play essential roles in diverse physiological and developmental processes, including transduction of environmental signals, establishing cell polarity, modulation of cellular motility, and regulating fluid flow (Pan, Wang et al. 2005, Berbari, O’Connor et al. 2009, Bloodgood 2010, Goetz and Anderson 2010, Dasgupta and Amack 2016, Ringers, Olstad et al. 2020). Dysfunction of cilia underlies a wide range of human syndromes—termed ciliopathies—that feature diverse phenotypes, including brain malformation, infertility, renal cyst formation, retinal degeneration, and anosmia (loss of smell) (Sharma, Berbari et al. 2008, Jenkins, McEwen et al. 2009, Brown and Witman 2014, Reiter and Leroux 2017, Uytingco, Green et al. 2019).

Cilia comprise three major compartments: the basal body (BB) with fibrous apparatuses transition fibers and basal feet, the transition zone (TZ), and the microtubule-based ciliary scaffold known as an “axoneme” (Kobayashi and Dynlacht 2011, Reiter, Blacque et al. 2012, Wei, Ling et al. 2015). Cilia are nucleated by the BB (derived from the mother centriole) and eventually protrude from the cell surface (Ishikawa and Marshall 2011). Subsequently, the TZ is templated to gate ciliary protein trafficking (Williams, Li et al. 2011). Known ciliary cargos include cilia structural components, G-protein-coupled receptors, ion channels, and other signaling molecules (Inglis, Ou et al. 2007, Lechtreck 2015, Nachury 2018). Those cargos are transported bi-directionally along the axoneme via a process called intraflagellar transport (IFT) (Hao and Scholey 2009). IFT components are recruited to elongate the ciliary axoneme. The IFT machinery consists of kinesin-2 and IFT-dynein motors, together with IFT-A and the IFT-B cargo adaptor complexes, that mediate the bidirectional movement of IFT cargos along the axoneme (Hao and Scholey 2009, Jordan and Pigino 2021). The anterograde IFT motors of the kinesin-2 family transport IFT particles from the cilia base to the cilia tip for incorporation into ciliary structures, while the retrograde IFT motors of dynein recycle kinesin-2 and IFT particles back to the cilia base (Hao and Scholey 2009, Prevo, Scholey et al. 2017). Besides, the Bardet Biedl syndrome (BBS) proteins are required to stabilize the association of IFT motors and IFT particles (Ou, Blacque et al. 2005, Uytingco, Williams et al. 2019). This bidirectional cargo transport is essential for ciliogenesis (Ishikawa and Marshall 2011). It has been reported that impairment of IFT leads to defects in cilia structure and function across different species. In *C. elegans*, mutations in IFT particle genes and motor genes have been shown to alter the cilia morphology of chemosensory neurons (Saikat Mukhopadhyay, Hongmin Qin et al. 2007). Inactivation of the IFT component IFT88 results in shortened cilia in a mouse model of polycystic kidney disease (Shao, El-Jouni et al. 2020). In addition, loss of BBS proteins leads to disorganization of the dendritic microtubule network of olfactory cilia, and causes anosmia in mice (Kulaga, Leitch et al. 2004).

Although the basic assembly mechanisms and structures of cilia are highly conserved, it is clear that these structures exhibit distinct lengths and morphologies depending on the identity and condition of the cells they are generated within; it is also clear that specialized cilia exert unique functional roles (Silverman and Leroux 2009). For example, the multiciliated protist model *Tetrahymena* carries two types of cilia (oral and locomotory) that exhibit asymmetries in the anterior-posterior and left-right axes (Soares, Carmona et al. 2019). These two types of cilia have different mechanisms to control cilia oscillation and to sense viscosity (Jung, Powers et al. 2014). In mammals, cilia of mammalian olfactory sensory neurons are known to have different lengths in distinct regions of the olfactory epithelium (Challis, Tian et al. 2015). Olfactory sensory neurons situated in the anterior areas have longer cilia and are more sensitive to odorants than those in the posterior regions (Challis, Tian et al. 2015). These findings make it clear that cilia identity (including morphology and function) is under strict control. However, any mechanisms through which the unique genesis, structural maintenance, and/or function of such cell-specific cilia remain elusive.

*C. elegans* has been repeatedly used as a model system to explore mechanisms regulating cell-specific cilia morphology and function (Inglis, Ou et al. 2007, Ou, Koga et al. 2007, Barr 2011). *C. elegans* has exactly 60 ciliated cells, with variable morphology and function (Bargmann 2006). Among the ciliated neurons, AWA, AWB, AWC, ASH, and ADL neurons belong to chemosensory neurons, enabling worms to detect a wide variety of volatile (olfactory) and water-soluble (gustatory) cues associated with food and danger (Emily R. Troemel 1997, Bargmann 2006, Hart and Chao 2010, Yoshida, Hirotsu et al. 2012, Li and Liberles 2015). Regarding morphology, AWA, AWB, and AWC neurons have “wing” cilia with distinct wing-like morphologies, while the ASH and ADL neurons have “channel” cilia with rod-like shapes (Inglis, Ou et al. 2007). For odorant recognition, the AWA and AWC neuron pairs detect volatile attractants (Cornelia I. Bargmann 1993, Sengupta 2007), while ASH, ADL, and AWB neurons respond to volatile repellants (Chao, Komatsu et al. 2004, Sengupta 2007). Thus, the cilia of particular chemosensory neurons of *C. elegans* represent an excellent model system to explore the cell-specific regulation of cilia morphology and function.

WD40-repeat (WDR) protein family is a large group of proteins containing the WDR motifs comprised of approximately 40 amino acids terminating in tryptophan (W) and aspartic acid (D) (Kim and Kim 2020). At least 17 different WDR proteins are associated with ciliopathies and majority of them have been identified as IFT components. One of the WDR proteins, WDR47 has been implicated in regulating formation of the central pair microtubules and ciliary beat in the motile cilia (Liu, Zheng et al. 2021). However, its function in the primary cilia, such as the chemosensory neurons remains unknow. Since WDR47 is highly conserved with NMTN-1 as the homolog in *C. elegans*, we intend to investigate if WDR47/NMTN-1 regulates the function of specific chemosensory neurons in *C. elegans*.

Here we discover that WDR47/NMTN-1 is required for AWB-mediated avoidance behaviors. After showing that WDR47/NMTN-1 is expressed in the AWB neuron pair and is enriched at the BB of AWB cilia, we demonstrate that knockout of *wdr47/nmtn-1* affects the cilia length and morphology of the AWB neurons as well as AWB-mediated chemosensation. Explaining these mutant phenotypes, our data support that WDR47/NMTN-1 helps to maintain appropriate IFT motor movement and proper IFT cargo delivery. In all, our results indicate that WDR47/NMTN-1 participates in the ciliogenesis via IFT particle movement in a cell-specific manner. Since WDR47/NMTN-1 is conserved in mammals, the mechanism we identified here may help us to better understand the process of cell-specific ciliogenesis and molecular mechanism for cilia identity.

## RESULTS

### WDR47/NMTN-1 is expressed in the AWB chemosensory neuron pair

To examine the expression pattern of WDR47/NMTN-1 in *C. elegans*, we generated transgenic animals expressing GFP under the *wdr47/nmtn-1* promoter. We detected the strong GFP signals in both the ciliated amphid and phasmid neurons (Fig. 1A). Next, we focused on the amphid chemosensory neurons and asked if WDR47/NMTN-1 is expressed in specific chemosensory neurons. There are five pairs of chemosensory neurons (olfactory) that detect volatile odors (Hart and Chao 2010). AWA and AWC neurons respond to volatile attractants (Cornelia I. Bargmann 1993, Sengupta 2007), while ASH, ADL, and AWB neurons respond to volatile repellants (Chao, Komatsu et al. 2004, Sengupta 2007). We labeled the individual chemosensory neuron pairs by expressing mCherry under neuron-type specific promoters (*odr-10* promoter for AWA, *str-1* promoter for AWB, *str-2* promoter for AWC, and *srb-6* promoter for ASH/ADL). We found strong GFP signals in the AWB neurons but not AWA or ASH/ADL neurons, and found a dim GFP signal in the AWC neurons (Fig. 1B-C). These data show that WDR47/NMTN-1 is expressed in the AWB chemosensory neuron pair known to function in chemosensation and aversion behaviors (Emily R. Troemel 1997).

**Figure 1.**
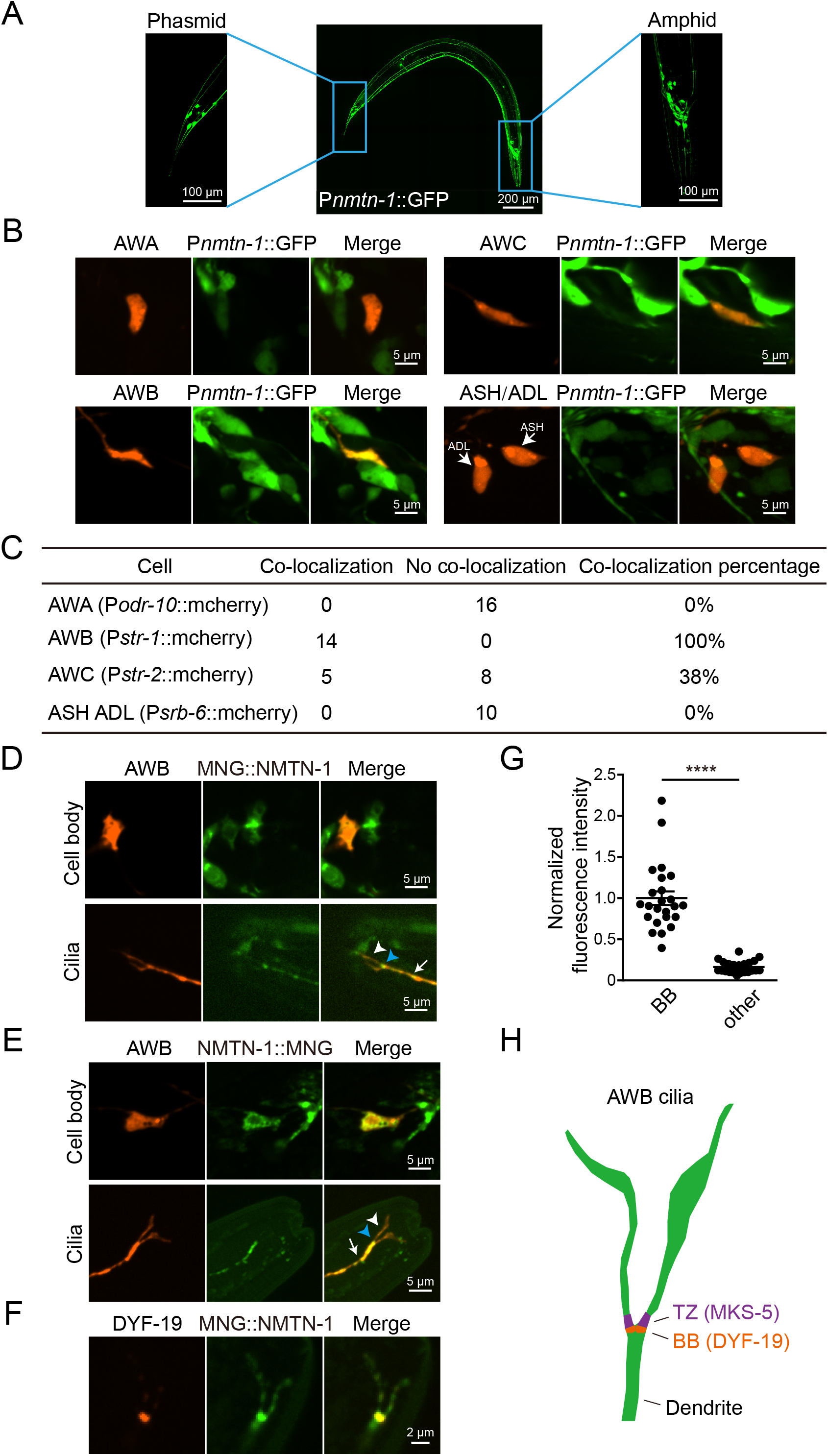
WDR47/NMTN-1 is expressed in the AWB chemosensory neuron pair and localized in the BB region. (A) WDR47/NMTN-1 is expressed in the amphids and phasmids of *C. elegans*. The left and right images are enlarged views of the phasmids and amphids, respectively. (B) Representative images of P*nmtn-1*::GFP signals in five pairs of olfactory neurons. The AWA, AWB, and AWC neurons are marked by P*odr-10*::mCherry, P*str-1*::mCherry, and P*str-2*::mCherry. The ASH and ADL neurons are marked by P*srb-6*::mCherry. (C) Quantification of P*nmtn-1*::GFP signals in five pairs of olfactory neurons. (D) Representative images showing P*nmtn-1*::MNG::NMTN-1 signals. The AWB neurons were visualized via expression of P*str-1*::mCherry. White arrowhead, blue arrowhead, and white arrow indicate cilia, the cilia base, and dendrites, respectively. (E) Representative images showing P*nmtn-1*::NMTN-1::MNG signals. The AWB neurons were marked by expression of P*str-1*::mCherry. White arrowhead, blue arrowhead, and white arrow indicate cilia, cilia base, and dendrites, respectively. (F) Colocalization of P*str-1*::MNG::NMTN-1 and the BB (P*str-1*::DYF-19::mCherry) marker. (G) Quantification of the fluorescence intensities of P*str-1*::MNG::NMTN-1 in BB and other parts of the cilia. Each cilium analyzed is represented by a dot. Data are presented as mean values ± SEM. **** P < 0.0001 by two-tailed Student’s t-test. (H) Schematic illustration of the AWB cilia and the location of transition zone (TZ) and basal body (BB). MKS-5 and DYF-19 are markers for the TZ and the BB, respectively.

### WDR47/NMTN-1 is localized at the basal body (BB) of cilia

To study the subcellular localization of WDR47/NMTN-1 in AWB neurons, we constructed a mNeonGreen-NMTN-1 (MNG::NMTN-1) fusion protein with expression under the *wdr47/nmtn-1* promoter (P*nmtn-1*). The mNeonGreen signals were enriched in the base of cilia and cell body of AWB neurons (Fig. 1D). We also constructed a C-terminal tagged NMTN-1-mNeonGreen (NMTN-1::MNG) fusion protein (same promoter), and found a similar localization pattern as the N-terminal tagged MNG::NMTN-1 (Fig. 1E). We also investigated the expression pattern of NMTN-1 at different developmental stages using the P*nmtn-1*::MNG::NMTN-1 fusion protein, and found that NMTN-1 was expressed in the cilia of the AWB neurons from egg to adult (day4) (Supplementary Fig. 1A-B).

There are two substructures at the base of cilia known to affect ciliogenesis and control ciliary protein composition: the basal body (BB) and the transition zone (TZ) (Fig. 1H) (Reiter, Blacque et al. 2012). Cilia are nucleated by the BB, and beyond the BB lies the TZ that acts as a “gate” to regulate the IFT-dependent trafficking of ciliary proteins to and from cilia (Ishikawa and Marshall 2011). To detect if WDR47/NMTN-1 is expressed in the BB and/or TZ, we labeled these substructures with mCherry-tagged MKS-5 and DYF-19 (Wei, Xu et al. 2013, Nechipurenko, Olivier-Mason et al. 2016). WDR47/NMTN-1 was co-localized with DYF-19 (Supplementary Fig. 2A-B), suggesting that WDR47/NMTN-1 is localized at the BB of the cilia of AWB neurons. We also verified the colocalization of WDR47/NMTN-1 and DYF-19 driven by the *str-1* promoter in the AWB neurons (Fig. 1F-G). The localization of WDR47/NMTN-1 in the BB implies that WDR47/NMTN-1 may regulate the BB structure. To test this possibility, we examined the distribution of mCherry-tagged DYF-19 proteins in *wdr47/nmtn-1* mutants. No abnormal localization of DYF-19 proteins was observed in *wdr47/nmtn-1* mutants, indicating that loss of WDR47/NMTN-1 may not affect the localization of ciliary base proteins (Supplementary Fig. 3).

### *Wdr47/nmtn-1* mutation causes morphology defects of AWB cilia

Given that the BB is known to function as a nucleation site for cilia biogenesis (Marshall 2008), we wondered whether WDR47/NMTN-1 might participate in the morphology of the AWB wing cilia. We classified the AWB cilia phenotypes into 3 categories using a previously reported method of quantifying AWB cilia morphologies (Olivier-Mason, Wojtyniak et al. 2013). Briefly, category 1 cilia have characteristic Y-shaped morphology with 2 primary branches and no fans. Category 2 cilia have enlarged fans along the primary branches. Category 3 cilia have more than one secondary branch emanating from the primary branch (Fig. 2A). We quantified the percentage of 3 categories in wild type and *wdr47/nmtn-1* mutants, and found that the percentage of category 1 cilia is significantly increased in *wdr47/nmtn-1* mutants, while the percentage of category 2 cilia is significantly decreased in these animals (Fig. 2B). We also measured the length of the AWB cilia: the typical AWB cilia contain 2 primary branches with unequal lengths, and we found that the lengths of both long and short branches in *wdr47/nmtn-1* mutants were significantly shorter than in wild type (Fig. 2C). These results collectively support that WDR47/NMTN-1 regulates cilia morphology of the AWB neurons.

**Figure 2.**
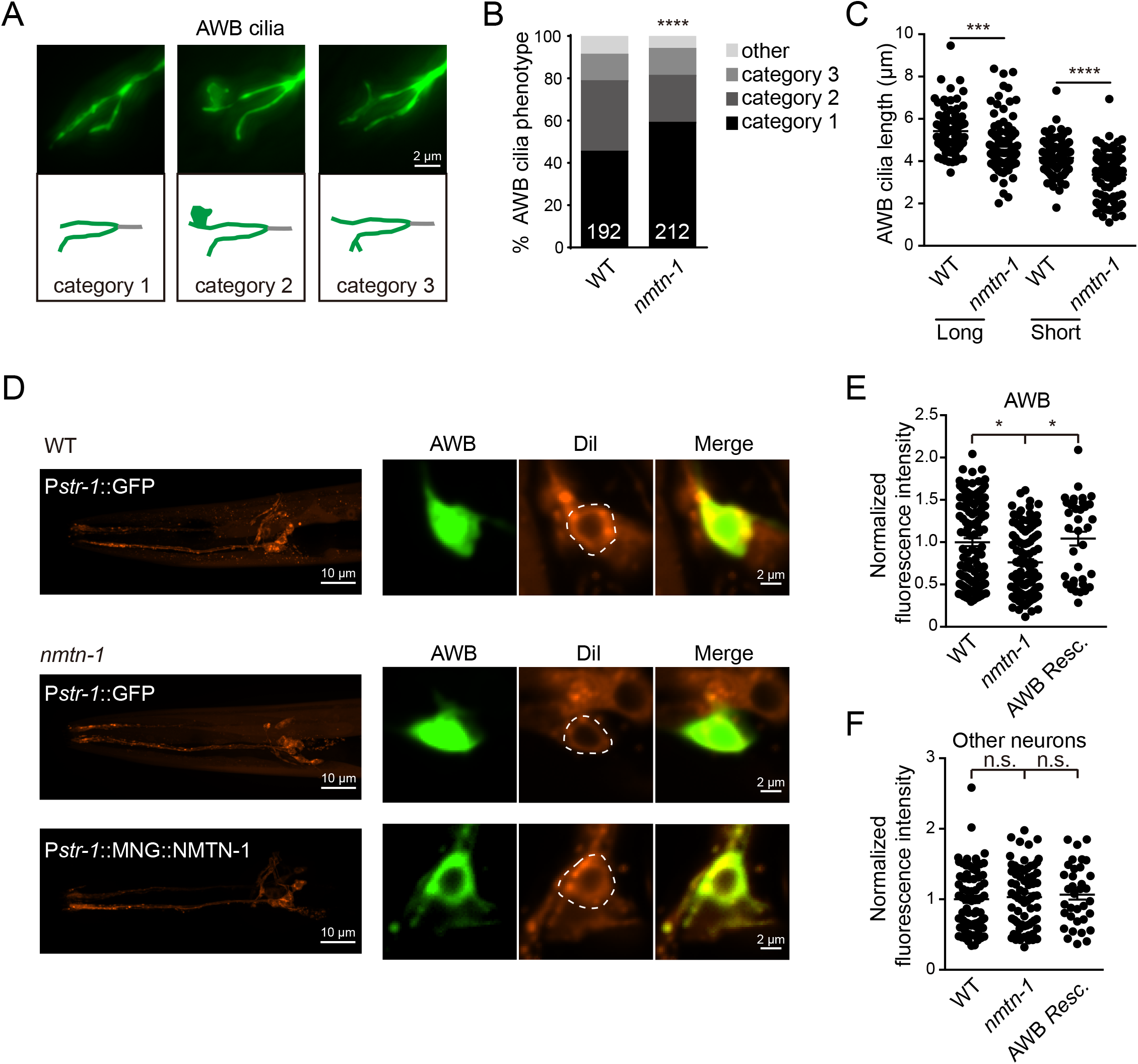
*Wdr47/nmtn-1* mutants exhibit defects in the AWB cilia morphology. (A) Representative images of the AWB cilia in three categories according to cilia morphology. Cilia were visualized using the P*str-1*::GFP marker, and were classified into three categories according to the cilia morphology. (B) Quantification of the AWB cilia in three categories in wild type (WT) and *wdr47/nmtn-1* mutants. (C) Quantification of the cilia length of the AWB neurons in WT and *wdr47/nmtn-1* mutants. Each cilium analyzed is represented by a dot. Data are presented as mean values ± SEM. (D-F) The *wdr47/nmtn-1* mutants exhibited cell-specific defects in uptake of the lipophilic dye Dil. Representative images of DiI uptake in amphid sensory neurons in WT and *wdr47/nmtn-1* mutants are displayed. The dotted lines represent AWB neurons (D). The dye-filling defect in AWB neurons of *wdr47/nmtn-1* animals was rescued by expression of P*str-1*::MNG::NMTN-1 (E). In contrast, the dye uptake in other neurons (random dye-filled non-AWB neurons) was normal in *wdr47/nmtn-1* mutants (F). Each neuron analyzed is represented by a dot. Data are presented as mean values ± SEM. In B, **** P < 0.0001 by Chi-square test. In C and E, * P < 0.05, *** P < 0.001, and **** P < 0.0001 by two-tailed Student’s t-test. In F, n.s. not significant by two-tailed Student’s t-test.

Recall our aforementioned observation of a dim WDR47/NMTN-1 signal in AWC neurons (Fig. 1B); we therefore examined AWC cilia morphology in *wdr47/nmtn-1* mutants. An abnormal morphology with discrete fan-shape cilia structure was occasionally observed in *wdr47/nmtn-1* mutants (16% in *wdr47/nmtn-1* mutants vs. 0% in wild type, Supplementary Fig. 4A-B). However, there were no significant differences in the AWC cilia area between *wdr47/nmtn-1* mutants and wild type animals (Supplementary Fig. 4C). We also investigated whether WDR47/NMTN-1 is required to maintain cilia morphology in other cell types using OSM-6-GFP fusion proteins to label cilia in all amphid ciliated sensory neurons. We found no significant changes in overall cilia length in *wdr47/nmtn-1* mutants in cells other than the AWB neurons (Supplementary Fig. 4D-E).

### *Wdr47/nmtn-1* mutation causes structural integrity defects of AWB cilia

We also performed a Dil dye-filling experiment—which is routinely used to validate the structural integrity of cilia (Tong and Burglin 2010)—to assess the specific impacts of WDR47/NMTN-1 in maintaining the AWB cilia structural integrity. After 30 minutes of Dil exposure, we analyzed the fluorescence intensity of Dil in the cell body. The dye signal in the AWB neurons of *wdr47/nmtn-1* mutants was significantly dimmer than in the wild type (Fig. 2D-E), suggesting apparent structural integrity defects in the mutant AWB cilia. In contrast, no Dil absorption defects were observed in other neurons (Fig. 2F). Furthermore, the Dil dye absorption defects in the AWB neurons of *wdr47/nmtn-1* mutants were restored upon the specific expression of WDR47/NMTN-1 in the AWB neurons driven by the *str-1* promoter (Fig. 2D-E). Those data suggest that WDR47/NMTN-1 functions to maintain the structural integrity of the AWB neuron cilia.

### WDR47/NMTN-1 is required for AWB-mediated aversion behaviors

The abnormal cilia morphology we detected in the AWB neurons of *wdr47/nmtn-1* mutants prompted us to test if WDR47/NMTN-1 is required for the AWB-mediated chemosensation behaviors. The AWB neurons are known to mediate aversion behaviors in response to odorants such as 2-nonanone, and these responses require intact and functional cilia (Emily R. Troemel 1997, Hart and Chao 2010). We conducted a classic chemotaxis assay to examine aversion behavior to 2-nonanone (Fig. 3A). Briefly, 9 cm plates containing regular NGM media were spotted with control (Ethanol) or 2-nonanone on opposite sides, and the paralysis agent sodium azide was added immediately before the addition of worms in the center of the plates (Fig. 3A). In line with the previous report (Cornelia I. Bargmann 1993), we observed that wild type animals were repelled by 2-nonanone and thus had a negative chemotaxis index value. The chemotaxis index value was significantly increased in the *wdr47/nmtn-1* mutants (Fig. 3B). Further, restoring WDR47/NMTN-1 expression either in the WDR47/NMTN-1-expressing neurons (under the *wdr47/nmtn-1* promoter) or in the AWB neurons (under the AWB-specific *str-1* promoter) rescued the chemotaxis defects (Fig. 3B). We also observed an impaired aversion response to octanol in the *wdr47/nmtn-1* mutants; octanol is known to act on a group of neurons including AWB (Fig. 3C) (Chao, Komatsu et al. 2004). No effects were observed when we investigated the attraction behaviors to different dilutions of diacetyl, mediated by the AWA neuron (Fig. 3D) (Cornelia I. Bargmann 1993). We did observe an impaired AWC-mediated attraction response to isopentyl alcohol in *wdr47/nmtn-1* mutants (Fig. 3E) (Cornelia I. Bargmann 1993), which may be due to the dim WDR47/NMTN-1 signal in the AWC neurons (Fig. 1B). Collectively, our data support the notion that WDR47/NMTN-1 functions in the AWB neurons to participate in the chemosensation behavior.

**Figure 3.**
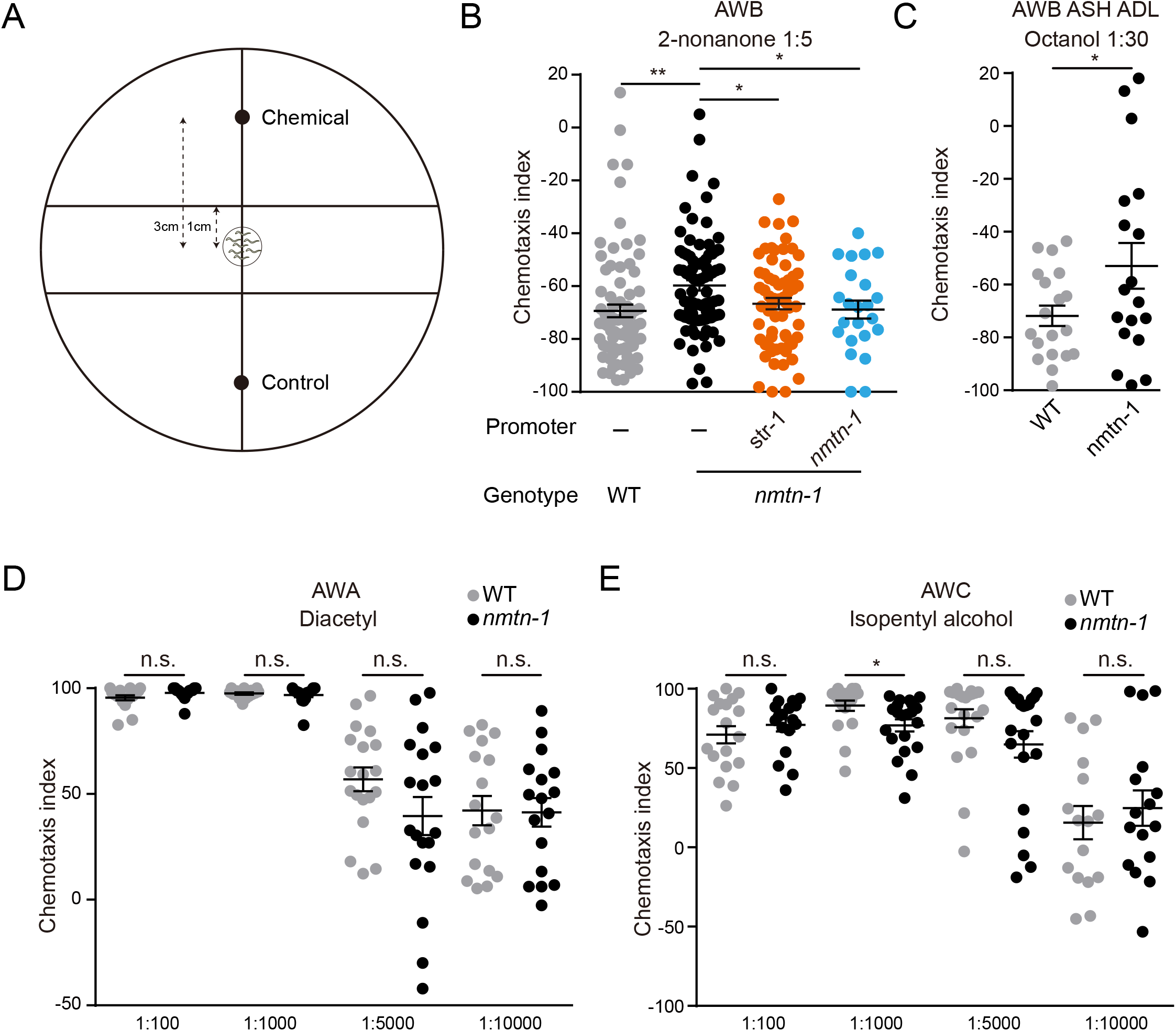
*Wdr47/nmtn-1* mutants have defects in AWB-mediated aversion behaviors. (A) Schematic illustration of the chemotaxis assays. (B) Quantification of chemotaxis indexes for AWB-mediated aversion behaviors to 2-nonanone in wild type (WT) and *wdr47/nmtn-1* mutants. The reduction in behavioral responses to 2-nonanone in *wdr47/nmtn-1* mutants can be rescued by expression of WDR47/NMTN-1 under its endogenous promoter or the AWB-specific *str-1* promoter. (C) Quantification of chemotaxis indexes for aversion behaviors to octanol mediated by the AWB, ASH, and ADL neurons in WT and *wdr47/nmtn-1* mutants. (D) Quantification of chemotaxis indexes for AWA-mediated attraction behaviors to diacetyl at the indicated concentrations in WT and *wdr47/nmtn-1* mutants. (E) Quantification of chemotaxis indexes for AWC-mediated attraction behaviors to isopentyl alcohol at the indicated concentrations in WT and *wdr47/nmtn-1* mutants. Each dot represents a single population assay calculated as shown. Data are presented as mean values ± SEM. In B, * P < 0.05, ** P < 0.01 by two-tailed Student’s t-test. In C and E, * P < 0.05, by two-tailed Student’s t-test. In D and E, n.s. not significant by two-tailed Student’s t-test.

### The overall ultrastructure of AWB cilia is not affected by *wdr47/nmtn-1* mutation

To explore the mechanisms underlying WDR47/NMTN-1’s impacts on ciliogenesis, we probed the ultrastructure of amphid cilia using transmission electron microscopy (TEM) (Serwas and Dammermann 2015). Unlike the axonemes of channel cilia, the microtubules in the AWB cilia lack an obvious organization (David B Doroquez, Berciu et al. 2014). We did not detect any microtubules in the distal segments of the AWB cilia, yet we did note the presence of singlet microtubules in the middle segments. However, no obvious abnormalities of the axoneme structure in the middle segments were observed in the *wdr47/nmtn-1* mutants (Supplementary Fig. 5), suggesting no disruption of the overall ultrastructure of AWB cilia.

### *Wdr47/nmtn-1* mutation perturbs the velocity distributions of IFT components

Ciliogenesis and cilia structure require the IFT-mediated bidirectional transport of particles along the microtubules (Hao and Scholey 2009), so we examined if deletion of *wdr47/nmtn-1* influences IFT particle movement. In *C. elegans*, two members of the kinesin-2 family, heterotrimeric kinesin-II (including KAP-1 protein) and homodimeric OSM-3 cooperate to form two sequential anterograde IFT pathways that build distinct parts of cilia (Scholey 2008). IFT particles involve two sub-complexes: IFT-A and IFT-B. IFT-A associates with kinesin-II, while IFT-B associates with OSM-3 during anterograde transport (Pedersen and Christensen 2012). To specifically examine IFT movement in the AWB neurons (Brust-Mascher, Ou et al. 2013), we expressed an MNG reporter fusion variant of KAP-1 and OSM-3 motor proteins, and the IFT-B complex subunit OSM-6 in the AWB neuron pair (under the AWB-specific *str-1* promoter) (Supplementary Fig. 6A). In the AWB cilia of wild type animals, the velocity of OSM-3 (0.87 μm/s) was a bit faster than that of KAP-1 and OSM-6 (0.64 μm/s and 0.63 μm/s) in the middle segments (Figure. 4A, Table 1). This is possibly due to the fact that some OSM-3 motors move alone in the middle segments, which is consistent with previous reports (Saikat Mukhopadhyay, Hongmin Qin et al. 2007). Interestingly, the velocity of OSM-3 (0.94 μm/s) was increased in the AWB cilia middle segments of *wdr47/nmtn-1* mutants, while the velocity of OSM-6 (0.53 μm/s) was decreased in *wdr47/nmtn-1* mutants (Fig. 4A, Table 1). We did not observe changes in the velocity of KAP-1. Those data indicate that loss of *wdr47/nmtn-1* perturbs the velocity distributions of IFT components. As controls, we did not detect significant changes in IFT velocities in the ASH or ADL cilia (under the *srb-6* promotor) (Table 1), further suggesting that WDR47/NMTN-1 is important for proper IFT movement in the AWB cilia.

**Table 1.** Transport velocities of MNG tagged IFT proteins in wild type (WT) and *wdr47/nmtn-1* mutant animals. n, number of particles; N, number of measured animals.

| IFT protein | Strain | Middle segment |  |  | Distal segment |  |  |
| --- | --- | --- | --- | --- | --- | --- | --- |
|  |  | Mean velocity (μm/s) | n/N | t test | Mean velocity (μm/s) | n/N | t test |
| AWB |  |  |  |  |  |  |  |
| *Pstr-1::KAP-1::MNG* | WT | 0.64 ± 0.01 | 81/5 |  | None |  |  |
| *Pstr-1::KAP-1::MNG* | *nmtn-1* | 0.64 ± 0.01 | 153/7 | 0.78 | None |  |  |
| *Pstr-1::OSM-3::MNG* | WT | 0.87 ± 0.02 | 104/7 |  | 1.14 ± 0.07 | 30/4 |  |
| *Pstr-1::OSM-3::MNG* | *nmtn-1* | 0.94 ± 0.01 | 160/4 | <0.001 | 1.16 ± 0.07 | 27/5 | 0.87 |
| *Pstr-1::OSM-6::MNG* | WT | 0.63 ± 0.02 | 70/5 |  | 1.11 ± 0.05 | 15/3 |  |
| *Pstr-1::OSM-6::MNG* | *nmtn-1* | 0.53 ± 0.01 | 80/7 | <0.001 | 0.79 ± 0.05 | 15/3 | <0.001 |
| ASH/ADL |  |  |  |  |  |  |  |
| *Psrb-6::OSM-6::MNG* | WT | 0.81 ± 0.01 | 180/8 |  | 1.39 ± 0.02 | 180/8 |  |
| *Psrb-6::OSM-6::MNG* | *nmtn-1* | 0.83 ± 0.01 | 175/6 | 0.17 | 1.37 ± 0.01 | 175/6 | 0.38 |

**Figure 4.**
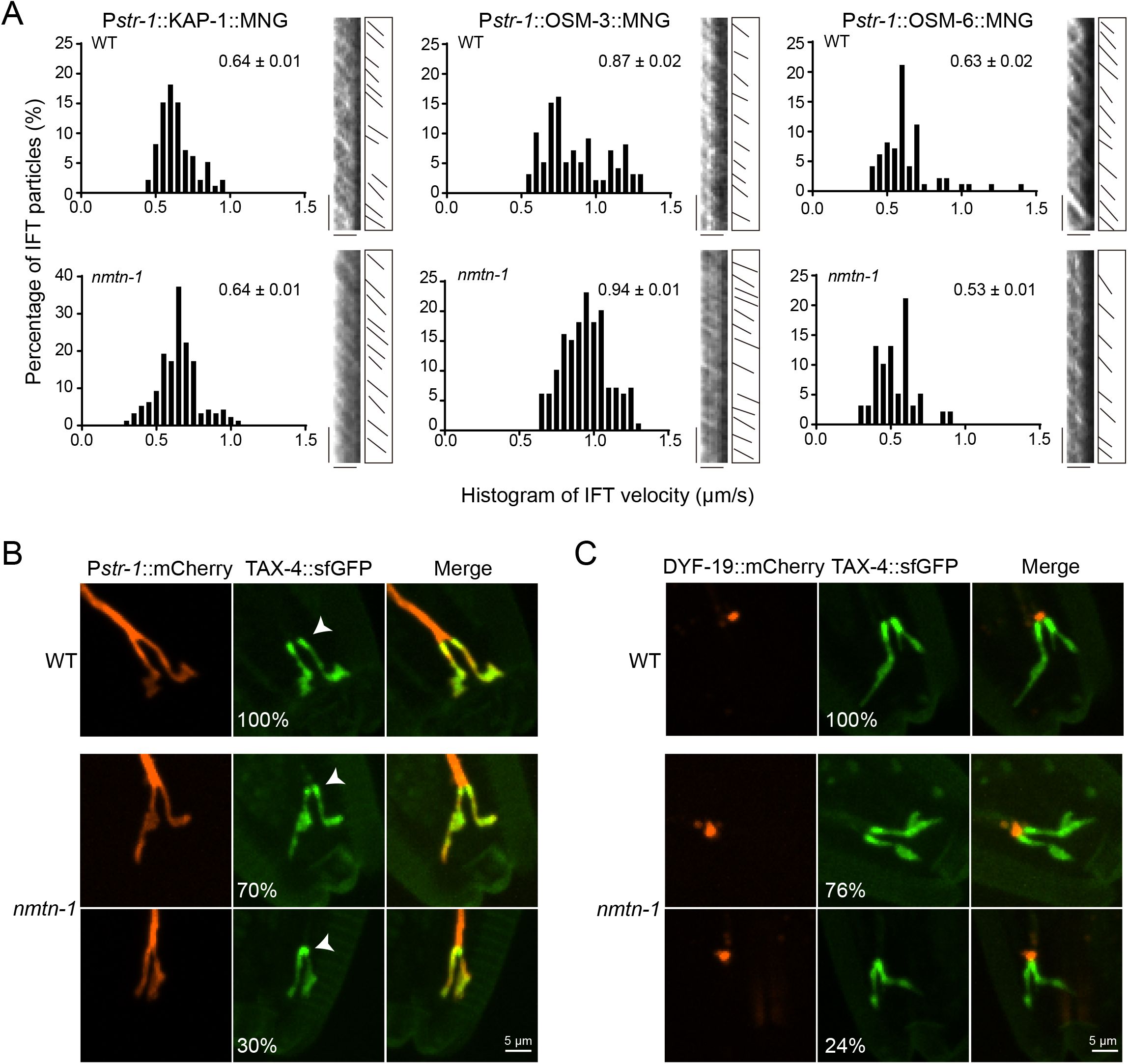
IFT velocities and localization of ciliary channel are altered in *wdr47/nmtn-1* mutants. (A) Histograms and kymographs of KAP-1::MNG, OSM-3::MNG, and OSM-6::MNG anterograde middle segment velocities in the AWB cilia of WT and *wdr47/nmtn-1* mutants. KAP-1::MNG, OSM-3::MNG and OSM-6::MNG were expressed under the AWB-specific *str-1* promoter. Average velocities are indicated at top right in each panel as mean values ± SEM. The scale bars represent 2 μm (horizontal) and 10 s (vertical). (B) Representative images of the P*str-1*::TAX-4::sfGFP fusion protein and the AWB cilia marked by P*str-1*::mCherry in WT and *wdr47/nmtn-1* mutants. TAX-4 is localized throughout cilia in all of WT animals, while TAX-4 is clearly detained the base of cilia in 30% of *wdr47****/****nmtn-1* mutants. (C) Representative images of the P*str-1*::TAX-4::sfGFP fusion protein and the BB protein marked by P*str-1*::DYF-19::mCherry in WT and *wdr47/nmtn-1* mutants. TAX-4 is localized throughout cilia in all of WT animals, while TAX-4 is clearly detained the base of cilia in 24% of *wdr47****/****nmtn-1* mutants. However, the detained TAX-4 in the base of AWB cilia is not co-localized with DYF-19.

To examine whether WDR47/NMTN-1 is one of the IFT components or otherwise physically associates with the IFT machinery, we analyzed the mobility of WDR47/NMTN-1 by kymograph: neither anterograde nor retrograde movement was observed (Supplementary Fig. 6B), indicating that the observed regulatory impacts of WDR47/NMTN-1 knockout on IFT movement may result from indirect interactions with IFT machinery.

### *Wdr47/nmtn-1* mutation alters IFT cargo localization

Since WDR47/NMTN-1 is required for IFT particle movement, we next examined whether IFT cargo transport requires WDR47/NMTN-1. As one of the IFT cargo, TAX-4 is the cyclic nucleotide-gated channel protein localized on the cilia membrane and participates in the olfactory signaling pathway (Bargmann 1996, Bargmann 2006). We studied the subciliary localization of TAX-4 in the *wdr47/nmtn-1* mutants. We expressed TAX-4::sfGFP fusion protein in the AWB neurons (under the *str-1* promoter) and found that TAX-4 was localized in the cilia in all of the wild type animals (Fig. 4B-C). In contrast, TAX-4 was mislocalized to the base of cilia in 20%-30% of *wdr47/nmtn-1* mutants (Fig. 4B). We also found that TAX-4 was mislocalized to the TZ region above the BB (Fig. 4C). These results further support the conclusion that WDR47/NMTN-1 is required for the IFT particle movement and cargo transportation, by which to support ciliogenesis and ciliary structure in the AWB neurons.

## DISCUSSION

In this study, we revealed how WDR47/NMTN-1 supports AWB cell-specific ciliogenesis and chemosensation in *C. elegans*. We showed that WDR47/NMTN-1 is expressed in the AWB chemosensory neurons and is enriched in the BB of the AWB cilia. WDR47/NMTN-1 functions in the AWB neurons to maintain AWB cilia morphology, structural integrity and AWB-mediated aversion behaviors. We further demonstrated that WDR47/NMTN-1 ensures proper IFT particle movement and cargo delivery in the AWB neurons, promoting ciliogenesis.

WDR47/NMTN-1 has been revealed as a microtubule-associated protein; it has been shown to interact with CAMSAP family proteins for microtubule-mediated processes (Chen, Zheng et al. 2020, Buijs, Hummel et al. 2021, Liu, Zheng et al. 2021). In non-centrosomal microtubules, WDR47/NMTN-1 protects CAMSAP2 against katanin-mediated severing and is required for axonal and dendritic development (Buijs, Hummel et al. 2021). In mammalian multicilia, WDR47/NMTN-1 co-operates with CAMSAP family proteins and MT-severing enzyme katanin to generate ciliary central microtubules (Liu, Zheng et al. 2021). WDR47/NMTN-1 also functions through CAMSAP3 to control neuronal migration and the early stages of neuronal polarization, which is important for neonatal mouse survival (Chen, Zheng et al. 2020). In addition, WDR47/NMTN-1 has been shown to interact with microtubule-associated protein 8 (Wang, Lundin et al. 2012) and participates in several microtubule-mediated processes including neural stem cell proliferation, radial migration, and growth cone dynamics (Kannan, Efil Bayam et al. 2017). Thus, multiple studies have conceptually linked WDR47/NMTN-1 with the regulation of microtubule-associated processes in neuron axons, dendrites, and motile cilia. In the present study, we discovered an additional role of WDR47/NMTN-1 in IFT particle movement and cell-specific ciliogenesis. It is likely that WDR47/NMTN-1 controls IFT particle movement via regulating ciliary microtubule networks.

Our results illustrate a cell-specific function of WDR47/NMTN-1 in ciliogenesis. This cell-specific modulation may have evolved to accommodate different olfactory receptors, channels, and/or IFT machinery in other chemosensory neurons to support diversified functions (Saikat Mukhopadhyay, Hongmin Qin et al. 2007, Silverman and Leroux 2009, Wojtyniak, Brear et al. 2013, 2014). Note that previous studies have reported that IFT-A molecules differentially regulate sensory cilia structures. IFT-121 and IFT-140 are required for all examined cilia in the amphid and phasmid neurons, whereas IFT-139 is required for ciliogenesis of AWC neuron-specific cilia (Scheidel and Blacque 2018). KLP-6, a conserved member of Kinesin-3 family, regulates IFT in the male-specific cilia (Morsci and Barr 2011). In addition, a few endocytic genes regulate ciliary and periciliary membrane compartment morphology in different cilia types, including the AWB cilia and 3 channel cilia (Kaplan, Doroquez et al. 2012). Similar to chemosensory neurons in *C. elegans*, mammalian olfactory sensory neurons are also divided into discrete subpopulations that contain distinct subfamilies of olfactory receptors in the cilia (Bear, Lassance et al. 2016). It will be quite interesting to explore the possibility that WDR47 orthologues may regulate the primary cilia of olfactory sensory neurons in mammals in a cell-specific manner.

Many WDR proteins, such as CHE-2/IFT80, WDR35/IFT121, DAF-10/IFT122, CHE-11/IFT140, and OSM-1/IFT172 are also required for ciliogenesis in analogy to WDR47/NMTN-1 (Manabi Fujiwara 1999, Qin, Rosenbaum et al. 2001, Quidwai, Wang et al. 2021). They are all mobile and act as IFT binding proteins. However, WDR47/NMTN-1 does not undergo IFT, so we speculate that WDR47/NMTN-1 may indirectly regulate IFT machinery. How does WDR47/NMTN-1 regulate the IFT velocities? Our observation that *wdr47/nmtn-1* perturbs the velocity distributions of IFT components has also been reported in *nphp-4* and *arl-13* mutants (Jauregui, Nguyen et al. 2008, Cevik, Hori et al. 2010). The velocity of OSM-3 is increased, while the velocity of OSM-6 is decreased in *nphp-4* and *arl-13* mutants. On the other hand, the velocity of KAP-1 is unchanged and decreased in *nphp-4* and *arl-13* mutants, respectively. They showed that OSM-6 is associated with kinesin-II other than OSM-3 in the absence of *nphp-4* (Jauregui, Nguyen et al. 2008), and OSM-3 is uncoupled from kinesin-II in *arl-13* mutants (Cevik, Hori et al. 2010). In wild type animals, the kinesin-II and OSM-3 units are linked by the BBS proteins, among these BBS-7 and BBS-8 are required to stabilize kinesin-II and OSM-3 (Ou, Blacque et al. 2005, Pan, Ou et al. 2006). Two kinases, the cell cycle-related kinase DYF-18 and the ros-cross hybridizing kinase family member MAK DYF-5 are important for stabilizing the interaction between IFT particles and OSM-3 (Yi, Xie et al. 2018). BBS components are predominantly localized at the base of cilia, and DYF-5 protein is mainly expressed in dendrites and TZ, and weakly expressed in cilia (Blacque, Reardon et al. 2004, Yi, Xie et al. 2018). Based on the fact that WDR47/NMTN-1 is localized in the BB of cilia, we suspect that WDR47/NMTN-1 may interact with BBS-7/8 and/or DYF-5/18 to maintain the coupling of OSM-3 and kinesin-II, or to regulate the binding of IFT particles with motor proteins. The overall effects lead to reduction of cargos transported to the cilia. This hypothesis is consistent with the observation that IFT cargo TAX-4 was detained in the base of the AWB cilia in some *wdr47/nmtn-1* mutants.

## MATERIALS AND METHODS

### Animals

*C. elegans* were maintained under standard conditions at 20 °C on nematode growth medium (NGM) plates seeded with *E. coli* OP50. All *C. elegans* strains were derived from the wild type Bristol N2 (Caenorhabditis Genetics Center) strain. The *wdr47/nmtn-1* mutant has a 483bp deletion in the second exon (chr1: 29916/29917-30399/30400). Transgenic animals were prepared by microinjection, and integrated transgenes were isolated following UV irradiation. A complete list of strains is provided in Supplementary Table 1.

### Plasmids

All expression vectors used are pPD49.26 or pPD95.75. A 3 kb *str-2* promoter was amplified from genomic DNA and cloned for expression in AWC chemosensory neurons. A 3 kb *odr-10* promoter was amplified from genomic DNA and cloned for expression in AWA chemosensory neurons. A 3 kb *str-1* promoter was amplified from genomic DNA and cloned for expression in AWB chemosensory neurons. A 3 kb *srb-6* promoter was amplified from genomic DNA and cloned for expression in ADF, ADL, and ASH chemosensory neurons. A complete list of primers used for cloning is provided in Supplementary Table 2.

### Live imaging and analysis

All hermaphrodites imaged were young adult animals. Worms were anesthetized with 30 µg/µl 2,3-butanedione monoxime (Sigma), mounted on the 2% agar pads. Fluorescent images were collected on a fluorescence microscope 100× (NA = 1.4) objective on an Olympus microscope (BX53) and a Nikon spinning disc confocal microscope (Yokogawa CSU-W1) equipped with a 60× oil objective. The images were further processed using ImageJ software.

For the time-lapse imaging experiment, worms were anesthetized with 10 mM levamisole, mounted on 5% agar pads. The images were taken on a Nikon spinning disc confocal microscope (Yokogawa CSU-W1) with a 60× oil objective. The exposure time of the time-lapse images is 300 ms. We used ImageJ software to process images, generate kymographs, and quantify IFT velocity. To ensure the quality of images used for quantification, only movies with worms in stable focal planes were used to generate kymographs. The anterograde kymographs were generated with the Reslice function in ImageJ by manually drawing lines along the AWB cilia.

### Dye-filling assay

Worms were washed with M9 buffer, and then incubated with the fluorescent lipophilic carbocyanine dye Dil in the dark for 30 min at room temperature. Dil was prepared as a 1 mg/ml stock solution in DMSO and diluted at 1:100 in M9 buffer. After incubation with Dil, worms were washed with M9 buffer again and transferred to seeded NGM plates for one or two hours to remove autofluorescence from the gut. Worms were then anesthetized with 10% levamisole, mounted on the 2% agar pads and imaged using a Nikon spinning disc confocal microscope (Yokogawa CSU-W1) with a 60× oil objective.

### Chemotaxis assay

The plates used for chemotaxis assay are 9 cm tissue culture dishes containing 10 ml of 1.6% agar, 5 mM potassium phosphate (pH 6.0), 1 mM CaCl_2_, and 1 mM MgSO_4_. The plates were autoclaved and stocked at 4°C. On the day of the experiments, the plates were taken out from 4°C to sit at room temperature until dry. The middle of the plates was marked on the back as the site for the initial location of the animals. In addition, two marks were labeled around 3 cm away from the middle of the plate. The two marks represent the sites for chemical and ethanol (control).

Synchronized young adult animals were washed three times with 1 ml S Basal buffer [5.9 g NaCl, 50 ml 1 M potassium phosphate (pH 6.0), 1 ml cholesterol (5 mg/ml in ethanol) in 1 L ddH_2_O]. Then the worms were washed two times with 1 ml water to remove bacteria, and were centrifuged at 3,000 rpm for 2 mins. The supernatants were removed as much as possible. 5 µl solution containing the worms was pipetted in the center of the plates. 1 µl of 1 M sodium azide was added to freeze animals on the control and chemical side. Next, 1 µl ethanol was added on the control side, and 1 µl chemical was added on the chemical side. After two hours, the number of animals on both the control and chemical side were counted. Note that the worms within 1 cm from the center were excluded. The chemotaxis indexes were calculated by using this formula: (Number of animals on the chemical side – Number of animals on control side) / (Number of animals on chemical side + Number of animals on the control side). The chemicals are provided in Supplementary Table 2.

### Transmission electron microscopy

We chose a type A carrier (100 µm + 200 µm, Leica, #16770181), dipped 200 µm surface with 1-hexadecene, and dried it with filter paper. The young adult hermaphrodites treated with 10 mM levamisole were transferred to M9 buffer containing 10% bovine serum albumin in the cavity of the carrier. The flat surface of the type B carrier (0 + 300 µm, Leica, #16770182) was placed on top to enclose the worms in the aluminum planchette’s cavity. The specimen–planchette sandwich was rapidly frozen using a Leica EM ICE high-pressure freezing system. Freeze-substitution was performed at low temperature (−90°C) over three days in a solution containing 2% osmium tetroxide, 1% uranyl acetate in anhydrous acetone using a Leica EM AFS2 freeze-substitution system. The temperature progressively increased up to 4°C. Samples were washed four times with anhydrous acetone (10 min each), and then successively infiltrated with a mixture of acetone resin of 3:1; 1:1; 1:3, respectively. Then samples were infiltrated and embedded in resin at room temperature and polymerized in an oven at 60°C for three days. Resin blocks with specimens were trimmed so that the block face was perpendicular to the longitudinal axis of the worm nose for sections, while keeping a small amount of resin around the specimen. Ultrathin sections (70 nm thickness) were collected and post-stained with 0.08 M lead citrate for 10 min. Sections were imaged on a 120 kV projection electron microscope (FEI, Talos L120C).

### Quantification and statistical analysis

All plots were generated by GraphPad Prism (version 7.0a). All scatterplots were shown as mean ± SEM. We used a two-tailed Student’s t-test to determine statistical differences except for the Chi-square test in Fig. 2B.

## ACKNOWLEDGEMENTS

We want to thank Xiumin Yan and Yidong Shen at the Institute of Biochemistry and Cell Biology, Shanghai Institutes for Biological Sciences, Chinese Academy of Sciences for kindly providing *wdr47/nmtn-1* mutants and fluorescent lipophilic carbocyanine dye Dil. We thank Guangshuo Ou at School of Life Sciences, Tsinghua University for kindly providing OSM-6::GFP strain and Qing Wei at Shenzhen Institutes of Advanced Technology, Chinese Academy of Sciences for providing P*arl13*::MKS-5::mCherry and P*arl13*::DYF-19::mCherry plasmids. This work was supported by the National Key Research and Development Program of China (2021ZD0203100), the National Natural Science Foundation of China (32122038, 31970933, and 32170963), the Basic Research Project from the Science and Technology Commission of Shanghai Municipality (21JC1404500 and 19JC1414100), Shuguang Program supported by Shanghai Education Development Foundation and Shanghai Municipal Education Commission (21SG16), Program for Young Scholars of Special Appointment at Shanghai Institutions of Higher Learning (QD2018017), and Innovative research team of high-level local universities in Shanghai (SHSMU-ZDCX20211102). We thank the *C. elegans* Genetics Stock Center, National BioResource Project (NBRP) for sharing strains and reagents. We also thank the Molecular Imaging Core Facility (MICF) at the School of Life Science and Technology, ShanghaiTech University for help in imaging.

## AUTHOR CONTRIBUTIONS

CS, XZ, WZ, XT, and QL designed experiments and analyzed data; CS, XT, and QL wrote the manuscript; XZ performed molecular cloning and microinjection experiments; WZ performed molecular cloning; CS performed molecular cloning, microinjection, imaging, and electron microscopy experiments.

## FIGURE LEGENDS

**Supplementary Figure 1. The expression pattern of P*nmtn-1*::MNG::NMTN-1 at different *C. elegans* developmental stages**. (A) Representative images showing P*nmtn-1*::MNG::NMTN-1 signals in the egg of *C. elegans*. The AWB neurons are marked by expression of P*str-1*::mCherry. (B) Representative images showing P*nmtn-1*::MNG::NMTN-1 signals in the L1, L2, L3, L4, day 1 adult, and day 4 adult of *C. elegans*. The AWB neurons were visualized by expression of P*str-1*::mCherry. The dotted lines represent the AWB neurons. White arrowhead, blue arrowhead, and white arrow indicate cilia, the cilia base, and dendrites, respectively.

**Supplementary Figure 2. Colocalization of P*str-1*::MNG::NMTN-1 with the TZ (MKS-5::mCherry) and BB (DYF-19::mCherry) markers**. (A-B) The arrowheads represent the positions of MKS-5 and DYF-19 in the AWB neurons. The dashed lines represent the cilia location. Fluorescence intensities are shown below each representative images. The dashed boxes show the regions for quantification of fluorescence intensity (starting from the lower right).

**Supplementary Figure 3. Representative images of P*str-1*::DYF-19::mCherry localization in the AWB neurons of wild type (WT) and *wdr47/nmtn-1* mutants**. AWB neurons were visualized using P*str-1*::mCherry.

**Supplementary Figure 4. *Wdr47/nmtn-1* mutants do not have defects in the morphology of other olfactory neurons**. (A) Representative images (top) and cartoons (bottom) of the normal and abnormal cilia of the AWC neurons. The AWC cilia were visualized by expression of P*str-2*::GFP. (B) The percentages of animals having the normal and abnormal cilia of the AWC neurons in wild type (WT) and *wdr47/nmtn-1* mutants are shown. (C) Quantification of the AWC cilia area in WT and *wdr47/nmtn-1* mutants. Each cilium analyzed is represented by a dot. Data are presented as mean values ± SEM. (D) Representative image of the OSM-6::GFP fusion protein. The line represents the length of cilia labeled by OSM-6::GFP. (E) Quantification of the cilia length labeled by OSM-6::GFP in WT and *wdr47/nmtn-1* mutants. Each cilium analyzed is represented by a dot. Data are presented as mean values ± SEM. In C and E, n.s. not significant by two-tailed Student’s t-test.

**Supplementary Figure 5. Ultrastructure of amphid cilia in wild type (WT) and *wdr47/nmtn-1* mutants**. Representative TEM images (cross-sections) of the amphid cilia distal and middle segment in WT and *wdr47/nmtn-1* mutants. White arrowheads indicate the AWB cilia.

**Supplementary Figure 6. Representative images of IFT movement**. (A) Representative images of P*str-1*::OSM-6::GFP, P*str-1*::OSM-3::GFP, P*str-1*::KAP-1::GFP, and P*srb-6*::OSM-6::GFP for analyses of IFT particle movement. (B) The kymograph image of WDR47/NMTN-1. The anterograde and retrograde kymograph images of P*nmtn-1*::MNG::NMTN-1 fusion protein show that WDR47/NMTN-1 is not moving by itself.

**Supplementary Table 1. List of *C. elegans* strains used in this study**. Summary of strain name, genotype, generating method, and resource of strains.

**Supplementary Table 2. List of chemicals, kits, and primers used for generating cell-specific promoters**.

## REFERENCES

Andrea G. Brear, J. Y., Martin Wojtyniak, and Piali Sengupta (2014). “Diverse Cell Type-Specific Mechanisms Localize G Protein-Coupled Receptors to Caenorhabditis elegans Sensory Cilia.” genetics 197: 667–684.

Bargmann, C. I. (2006). “Chemosensation in C. elegans.” WormBook: 1–29.

Bargmann, C. M. C. a. C. I. (1996). “A Putative Cyclic Nucleotide–Gated Channel.” Neuron 17: 695–706.

Barr, Y.-K. B. a. M. M. (2011). “Sensory roles of neuronal cilia: Cilia development, morphogenesis, and function in C. elegans.” Front Biosci. 13: 5959–5974.

Bear, D. M., J. M. Lassance, H. E. Hoekstra and S. R. Datta (2016). “The Evolving Neural and Genetic Architecture of Vertebrate Olfaction.” Curr Biol 26(20): R1039–R1049.

Berbari, N. F., A. K. O’Connor, C. J. Haycraft and B. K. Yoder (2009). “The primary cilium as a complex signaling center.” Curr Biol 19(13): R526–535.

Blacque, O. E., M. J. Reardon, C. M. Li, J. McCarthy, M. R. Mahjoub, S. J. Ansley, L. L. Badano, A. K. Mah, P. L. Beales, W. S. Davidson, R. C. Johnsen, M. Audeh, R. H. A. Plasterk, D. L. Baillie, N. Katsanis, L. M. Quarmby, S. R. Wicks and M. R. Leroux (2004). “Loss of C-elegans BBS-7 and BBS-8 protein function results in cilia defects and compromised intraflagellar transport.” Genes & Development 18(13): 1630–1642.

Bloodgood, R. A. (2010). “Sensory reception is an attribute of both primary cilia and motile cilia.” J Cell Sci 123(Pt 4): 505–509.

Brown, J. M. and G. B. Witman (2014). “Cilia and Diseases.” Bioscience 64(12): 1126–1137.

Brust-Mascher, I., G. Ou and J. M. Scholey (2013). “Measuring rates of intraflagellar transport along Caenorhabditis elegans sensory cilia using fluorescence microscopy.” Methods Enzymol 524: 285–304.

Buijs, R. R., J. J. A. Hummel, M. Burute, X. Pan, Y. Cao, R. Stucchi, M. Altelaar, A. Akhmanova, L. C. Kapitein and C. C. Hoogenraad (2021). “WDR47 protects neuronal microtubule minus ends from katanin-mediated severing.” Cell Rep 36(2): 109371.

Cevik, S., Y. Hori, O. I. Kaplan, K. Kida, T. Toivenon, C. Foley-Fisher, D. Cottell, T. Katada, K. Kontani and O. E. Blacque (2010). “Joubert syndrome Arl13b functions at ciliary membranes and stabilizes protein transport in Caenorhabditis elegans.” J Cell Biol 188(6): 953–969.

Challis, R. C., H. Tian, J. Wang, J. He, J. Jiang, X. Chen, W. Yin, T. Connelly, L. Ma, C. R. Yu, J. L. Pluznick, D. R. Storm, L. Huang, K. Zhao and M. Ma (2015). “An Olfactory Cilia Pattern in the Mammalian Nose Ensures High Sensitivity to Odors.” Curr Biol 25(19): 2503–2512.

Chao, M. Y., H. Komatsu, H. S. Fukuto, H. M. Dionne and A. C. Hart (2004). “Feeding status and serotonin rapidly and reversibly modulate a Caenorhabditis elegans chemosensory circuit.” Proc Natl Acad Sci U S A 101(43): 15512–15517.

Chen, Y., J. Zheng, X. Li, L. Zhu, Z. Shao, X. Yan and X. Zhu (2020). “Wdr47 Controls Neuronal Polarization through the Camsap Family Microtubule Minus-End-Binding Proteins.” Cell Rep 31(3): 107526.

Cornelia I. Bargmann, t. E. H., and H. Robert Horvitz (1993). “Odorant-Selective Genes and Neurons Mediate Olfaction in C. elegans.” Cell 74: 515–527,.

Dasgupta, A. and J. D. Amack (2016). “Cilia in vertebrate left-right patterning.” Philos Trans R Soc Lond B Biol Sci 371(1710).

David B Doroquez, C. Berciu and D. Nicastro (2014). “A high-resolution morphological and ultrastructural map of anterior sensory cilia and glia in Caenorhabditis elegans.” eLife.

Emily R. Troemel, B. E. K., and Cornelia I. Bargmann (1997). “Reprogramming Chemotaxis Responses: Sensory Neurons Define Olfactory Preferences in C. elegans.” Cell 91: 161–169.

Goetz, S. C. and K. V. Anderson (2010). “The primary cilium: a signalling centre during vertebrate development.” Nat Rev Genet 11(5): 331–344.

Hao, L. and J. M. Scholey (2009). “Intraflagellar transport at a glance.” Cell Science at a Glance 122: 889–892.

Hart, A. C. and M. Y. Chao (2010). Frontiers in Neuroscience.From Odors to Behaviors in Caenorhabditis elegans. The Neurobiology of Olfaction.

Inglis, P. N., G. Ou, M. R. Leroux and J. M. Scholey (2007). “The sensory cilia of Caenorhabditis elegans.” WormBook: 1–22.

Ishikawa, H. and W. F. Marshall (2011). “Ciliogenesis: building the cell’s antenna.” Nat Rev Mol Cell Biol 12(4): 222–234.

Jauregui, A. R., K. C. Q. Nguyen, D. H. Hall and M. M. Barr (2008). “The Caenorhabditis elegans nephrocystins act as global modifiers of cilium structure.” J Cell Biol 180(5): 973–988.

Jenkins, P. M., D. P. McEwen and J. R. Martens (2009). “Olfactory cilia: linking sensory cilia function and human disease.” Chem Senses 34(5): 451–464.

Jordan, M. A. and G. Pigino (2021). “The structural basis of intraflagellar transport at a glance.” J Cell Sci 134(12).

Jung, I., T. R. Powers and J. M. Valles, Jr. (2014). “Evidence for two extremes of ciliary motor response in a single swimming microorganism.” Biophys J 106(1): 106–113.

Kannan, M., Efil Bayam, C. Wagner, Bruno Rinaldif, Perrine F. Kretz, P. Tilly, Marna Roosg, Lara McGillewieh, Séverine Bärf, Shilpi Minochae, Claire Chevalier, Chrystelle Poi, S. M. G. Project, J. Chelly, J.-L. Mandel, Renato Borgattik, A. Piton, Craig Kinnearh, Ben Loosg, D. J. Adams, Y. Hérault, S. C. Collins, Sylvie Friantf, J. D. Godin and B. Yalcin (2017). “WD40-repeat 47, a microtubule-associated protein, is essential for brain development and autophagy.” PNAS: E9308–E9317.

Kaplan, O. I., D. B. Doroquez, S. Cevik, R. V. Bowie, L. Clarke, A. A. Sanders, K. Kida, J. Z. Rappoport, P. Sengupta and O. E. Blacque (2012). “Endocytosis genes facilitate protein and membrane transport in C. elegans sensory cilia.” Curr Biol.

Kim, Y. and S. H. Kim (2020). “WD40-Repeat Proteins in Ciliopathies and Congenital Disorders of Endocrine System.” Endocrinol Metab (Seoul) 35(3): 494–506.

Kobayashi, T. and B. D. Dynlacht (2011). “Regulating the transition from centriole to basal body.” J Cell Biol 193(3): 435–444.

Kulaga, H. M., C. C. Leitch, E. R. Eichers, J. L. Badano, A. Lesemann, B. E. Hoskins, J. R. Lupski, P. L. Beales, R. R. Reed and N. Katsanis (2004). “Loss of BBS proteins causes anosmia in humans and defects in olfactory cilia structure and function in the mouse.” Nat Genet 36(9): 994–998.

Lechtreck, K. F. (2015). “IFT-Cargo Interactions and Protein Transport in Cilia.” Trends Biochem Sci 40(12): 765–778.

Li, Q. and S. D. Liberles (2015). “Aversion and attraction through olfaction.” Curr Biol 25(3): R120–129.

Liu, H., J. Zheng, L. Zhu, L. Xie, Y. Chen, Y. Zhang, W. Zhang, Y. Yin, C. Peng, J. Zhou, X. Zhu and X. Yan (2021). “Wdr47, Camsaps, and Katanin cooperate to generate ciliary central microtubules.” Nat Commun 12(1): 5796.

Manabi Fujiwara, T. I. a. I. K. (1999). “A novel WD40 protein, CHE-2, acts cell autonomously in the formation of C. elegans sensory cilia.” Development(126): 4839–4848

Marshall, W. F. (2008). Chapter 1 Basal Bodies. Ciliary Function in Mammalian Development: 1–22.

Morsci, N. S. and M. M. Barr (2011). “Kinesin-3 KLP-6 regulates intraflagellar transport in male-specific cilia of Caenorhabditis elegans.” Curr Biol 21(14): 1239–1244.

Nachury, M. V. (2018). “The molecular machines that traffic signaling receptors into and out of cilia.” Curr Opin Cell Biol 51: 124–131.

Nechipurenko, I. V., A. Olivier-Mason, A. Kazatskaya, J. Kennedy, I. G. McLachlan, M. G. Heiman, O. E. Blacque and P. Sengupta (2016). “A Conserved Role for Girdin in Basal Body Positioning and Ciliogenesis.” Dev Cell 38(5): 493–506.

Olivier-Mason, A., M. Wojtyniak, R. V. Bowie, I. V. Nechipurenko, O. E. Blacque and P. Sengupta (2013). “Transmembrane protein OSTA-1 shapes sensory cilia morphology via regulation of intracellular membrane trafficking in C. elegans.” Development 140(7): 1560–1572.

Ou, G., O. E. Blacque, J. J. Snow, M. R. Leroux and J. M. Scholey (2005). “Functional coordination of intraflagellar transport motors.” Nature 436(7050): 583–587.

Ou, G., M. Koga, O. E. Blacque, T. Murayama, Y. Ohshima, J. C. Schafer, C. Li, B. K. Yoder, M. R. Leroux and J. M. Scholey (2007). “Sensory ciliogenesis in Caenorhabditis elegans: assignment of IFT components into distinct modules based on transport and phenotypic profiles.” Mol Biol Cell 18(5): 1554–1569.

Pan, J., Q. Wang and W. J. Snell (2005). “Cilium-generated signaling and cilia-related disorders.” Lab Invest 85(4): 452–463.

Pan, X., G. Ou, G. Civelekoglu-Scholey, O. E. Blacque, N. F. Endres, L. Tao, A. Mogilner, M. R. Leroux, R. D. Vale and J. M. Scholey (2006). “Mechanism of transport of IFT particles in C. elegans cilia by the concerted action of kinesin-II and OSM-3 motors.” J Cell Biol 174(7): 1035–1045.

Pedersen, L. B. and S. T. Christensen (2012). “Regulating intraflagellar transport.” Nat Cell Biol 14(9): 904–906.

Pedersen, L. B., J. M. Schroder, P. Satir and S. T. Christensen (2012). “The ciliary cytoskeleton.” Compr Physiol 2(1): 779–803.

Prevo, B., J. M. Scholey and E. J. G. Peterman (2017). “Intraflagellar transport: mechanisms of motor action, cooperation, and cargo delivery.” FEBS J 284(18): 2905–2931.

Qin, H. M., J. L. Rosenbaum and M. M. Barr (2001). “An autosomal recessive polycystic kidney disease gene homolog is involved in intraflagellar transport in C-elegans ciliated sensory neurons.” Current Biology 11(6): 457–461.

Quidwai, T., J. Wang, E. A. Hall, N. A. Petriman, W. Leng, P. Kiesel, J. N. Wells, L. C. Murphy, M. A. Keighren, J. A. Marsh, E. Lorentzen, G. Pigino and P. Mill (2021). “A WDR35-dependent coat protein complex transports ciliary membrane cargo vesicles to cilia.” Elife 10.

Reiter, J. F., O. E. Blacque and M. R. Leroux (2012). “The base of the cilium: roles for transition fibres and the transition zone in ciliary formation, maintenance and compartmentalization.” EMBO Rep 13(7): 608–618.

Reiter, J. F. and M. R. Leroux (2017). “Genes and molecular pathways underpinning ciliopathies.” Nat Rev Mol Cell Biol 18(9): 533–547.

Ringers, C., E. W. Olstad and N. Jurisch-Yaksi (2020). “The role of motile cilia in the development and physiology of the nervous system.” Philos Trans R Soc Lond B Biol Sci 375(1792): 20190156.

Saikat Mukhopadhyay, Y. L., A. L. Hongmin Qin and S. S. a. P. Sengupta (2007). “Distinct IFT mechanisms contribute to the generation of ciliary structural diversity in C. elegans.” The EMBO Journal 26: 2966–2980.

Scheidel, N. and O. E. Blacque (2018). “Intraflagellar Transport Complex A Genes Differentially Regulate Cilium Formation and Transition Zone Gating.” Curr Biol 28(20): 3279-3287.e3272.

Scholey, J. M. (2008). “Intraflagellar transport motors in cilia: moving along the cell’s antenna.” J Cell Biol 180(1): 23–29.

Sengupta, P. (2007). “Generation and modulation of chemosensory behaviors in C. elegans.” Pflugers Arch 454(5): 721–734.

Serwas, D. and A. Dammermann (2015). “Ultrastructural analysis of Caenorhabditis elegans cilia.” Methods Cell Biol 129: 341–367.

Shao, L., W. El-Jouni, F. Kong, J. Ramesh, R. S. Kumar, X. Shen, J. Ren, S. Devendra, A. Dorschel, M. Wu, I. Barrera, A. Tabari, K. Hu, N. Haque, I. Yambayev, S. Li, A. Kumar, T. R. Behera, G. McDonough, M. Furuichi, M. Xifaras, T. Lu, R. M. Alhayaza, K. Miyabayashi, Q. Fan, A. K. Ajay and J. Zhou (2020). “Genetic reduction of cilium length by targeting intraflagellar transport 88 protein impedes kidney and liver cyst formation in mouse models of autosomal polycystic kidney disease.” Kidney Int 98(5): 1225–1241.

Sharma, N., N. F. Berbari and B. K. Yoder (2008). “Ciliary dysfunction in developmental abnormalities and diseases.” Curr Top Dev Biol 85: 371–427.

Silverman, M. A. and M. R. Leroux (2009). “Intraflagellar transport and the generation of dynamic, structurally and functionally diverse cilia.” Trends Cell Biol 19(7): 306–316.

Soares, H., B. Carmona, S. Nolasco and L. Viseu Melo (2019). “Polarity in Ciliate Models: From Cilia to Cell Architecture.” Front Cell Dev Biol 7: 240.

Tong, Y. G. and T. R. Burglin (2010). “Conditions for dye-filling of sensory neurons in Caenorhabditis elegans.” J Neurosci Methods 188(1): 58–61.

Uytingco, C. R., W. W. Green and J. R. Martens (2019). “Olfactory Loss and Dysfunction in Ciliopathies: Molecular Mechanisms and Potential Therapies.” Curr Med Chem 26(17): 3103–3119.

Uytingco, C. R., C. L. Williams, C. Xie, D. T. Shively, W. W. Green, K. Ukhanov, L. Zhang, D. Y. Nishimura, V. C. Sheffield and J. R. Martens (2019). “BBS4 is required for intraflagellar transport coordination and basal body number in mammalian olfactory cilia.” J Cell Sci 132(5).

Wang, W., V. F. Lundin, I. Millan, A. Zeng, X. Chen, J. Yang, E. Allen, N. Chen, G. Bach, A. Hsu, M. T. Maloney, M. Kapur and Y. Yang (2012). “Nemitin, a novel Map8/Map1s interacting protein with Wd40 repeats.” PLoS One 7(4): e33094.

Wei, Q., K. Ling and J. Hu (2015). “The essential roles of transition fibers in the context of cilia.” Curr Opin Cell Biol 35: 98–105.

Wei, Q., Q. Xu, Y. Zhang, Y. Li, Q. Zhang, Z. Hu, P. C. Harris, V. E. Torres, K. Ling and J. Hu (2013). “Transition fibre protein FBF1 is required for the ciliary entry of assembled intraflagellar transport complexes.” Nat Commun 4: 2750.

Williams, C. L., C. Li, K. Kida, P. N. Inglis, S. Mohan, L. Semenec, N. J. Bialas, R. M. Stupay, N. Chen, O. E. Blacque, B. K. Yoder and M. R. Leroux (2011). “MKS and NPHP modules cooperate to establish basal body/transition zone membrane associations and ciliary gate function during ciliogenesis.” J Cell Biol 192(6): 1023–1041.

Wojtyniak, M., A. G. Brear, D. M. O’Halloran and P. Sengupta (2013). “Cell-and subunit-specific mechanisms of CNG channel ciliary trafficking and localization in C. elegans.” J Cell Sci 126(Pt 19): 4381–4395.

Yi, P., C. Xie and G. Ou (2018). “The kinases male germ cell-associated kinase and cell cycle-related kinase regulate kinesin-2 motility in Caenorhabditis elegans neuronal cilia.” Traffic 19(7): 522–535.

Yoshida, K., T. Hirotsu, T. Tagawa, S. Oda, T. Wakabayashi, Y. Iino and T. Ishihara (2012). “Odour concentration-dependent olfactory preference change in C. elegans.” Nat Commun 3: 739.

