## Supplementary figures and images for "WDR47 facilitates ciliogenesis by modulating intraflagellar transport"

### Supplementary Figure 1

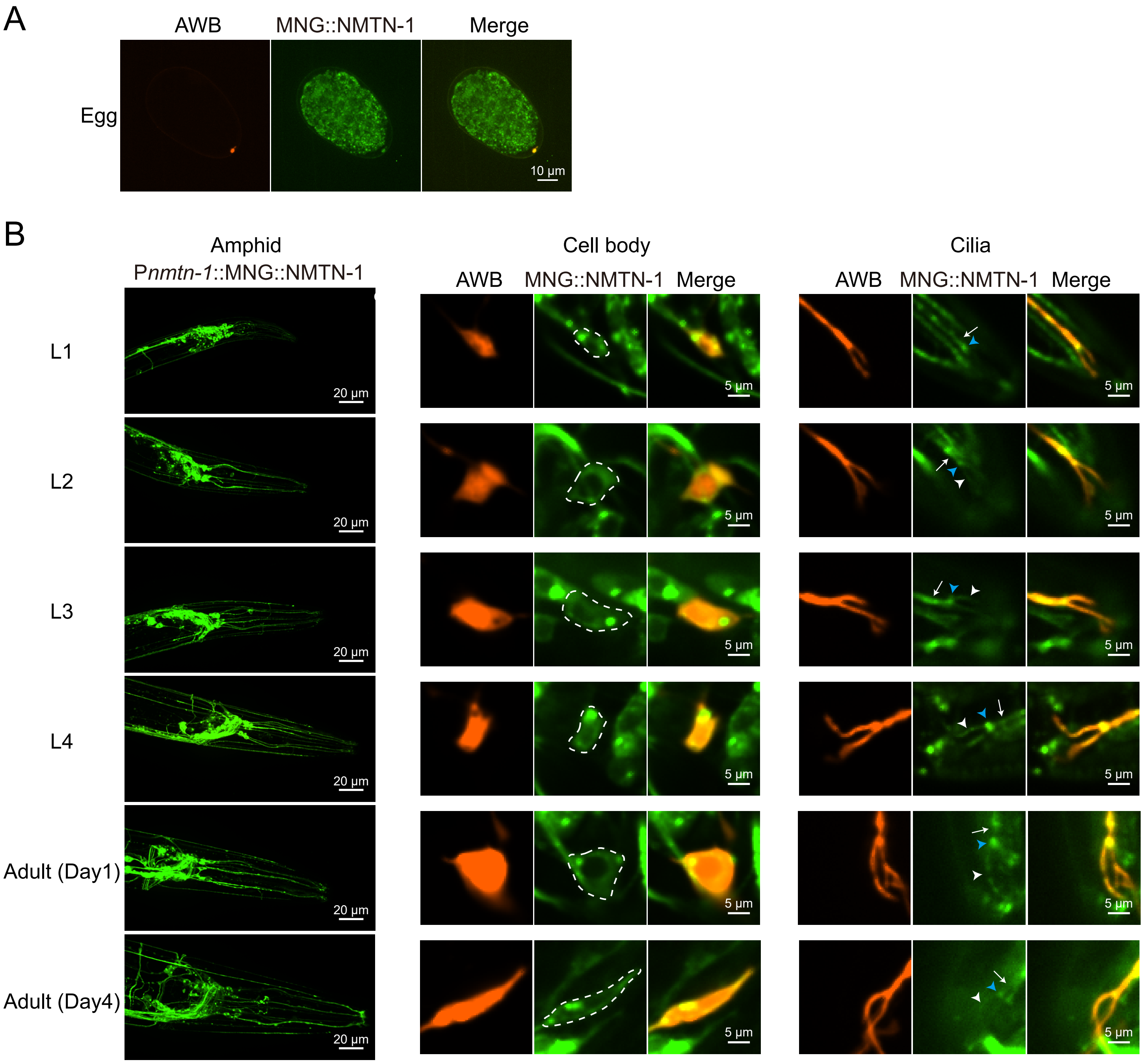

### Supplementary Figure 2

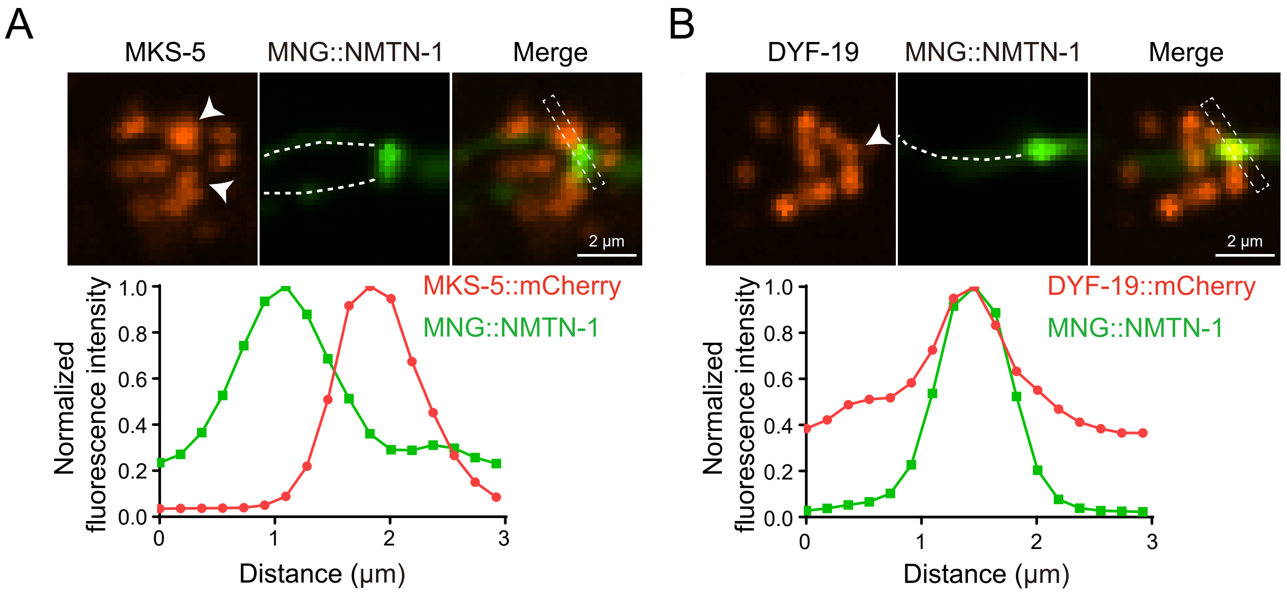

### Supplementary Figure 3

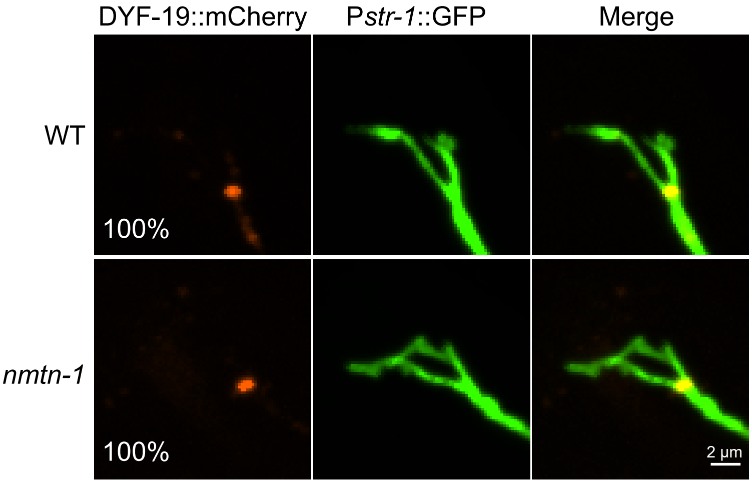


Figure S3

### Supplementary Figure 4

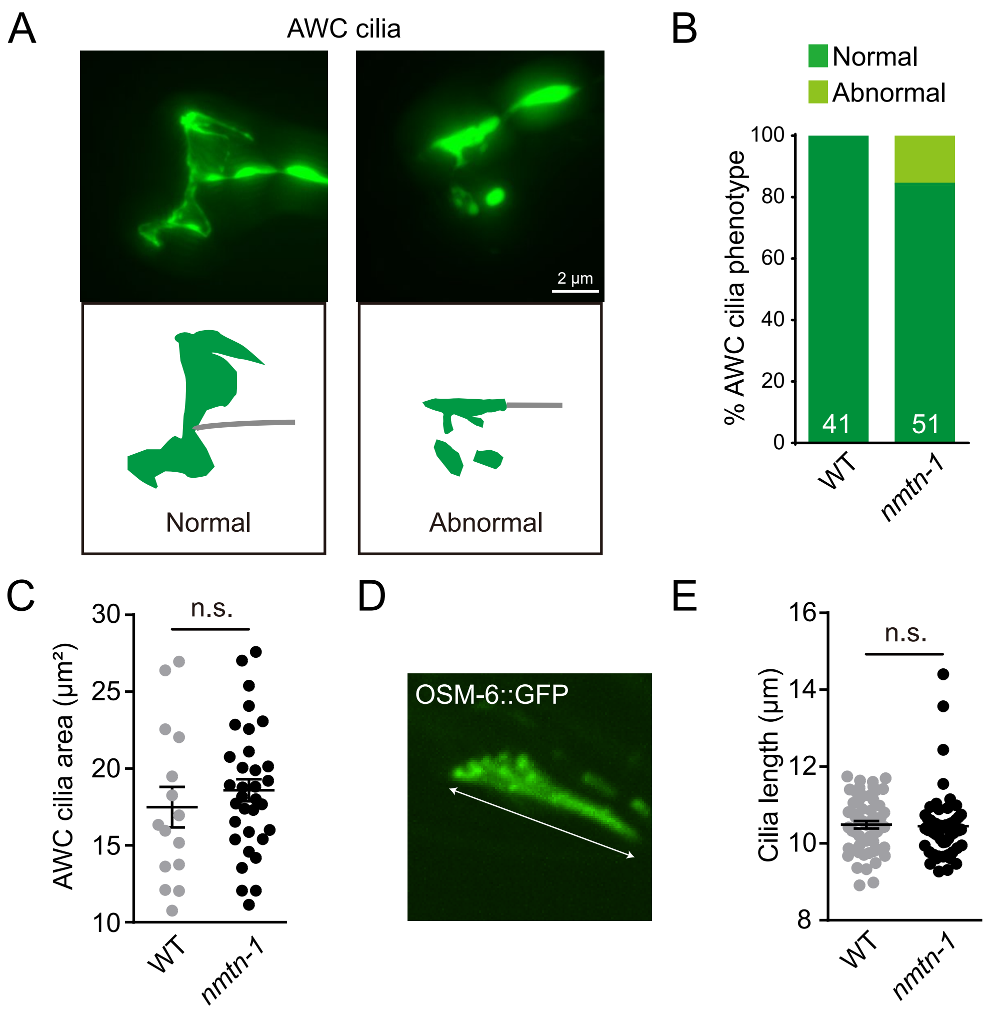

### Supplementary Figure 5

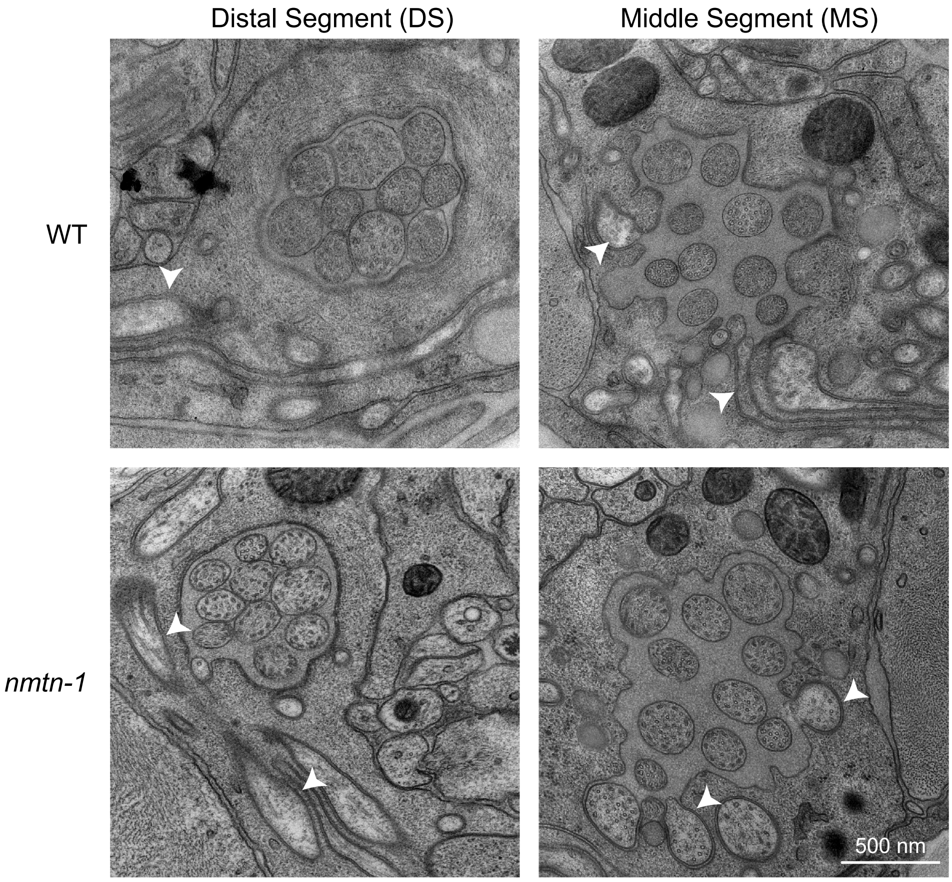


Figure S5

### Supplementary Figure 6

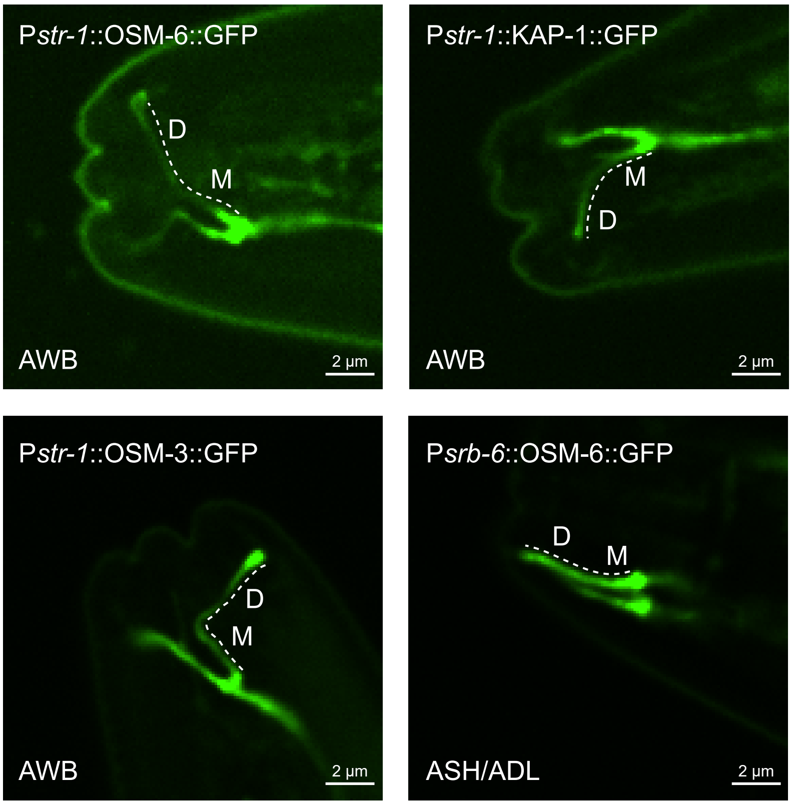

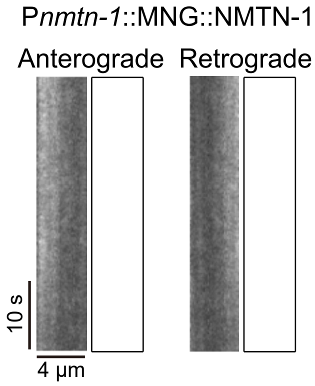
