## Supplementary Table 1 for "WDR47 facilitates ciliogenesis by modulating intraflagellar transport"

| Supplementary Table 1. *C. elegans* strains used in this study | | | |
|---|---|---|---|
| **Strain name** | **Genotype** | **Method** | **Resource** |
| N2 | Wild type | - | CGC |
| TM5438 | *nmtn-1(tm5438) I* | - | NBRP |
| TXJ0539 | *xjIs0003[Pstr-1::GFP+Pmyo-3:: mCherry]* | Microinjection | In this study |
| TXJ0553 | *nmtn-1(tm5438) I; xjIs0003[Pstr-1::GFP+Pmyo-3:: mCherry]* | Genetic cross | In this study |
| SYD0202 | *[OSM-6::GFP]* | - | Guangshuo Ou |
| TXJ0649 | *nmtn-1(tm5438) I;[OSM-6::GFP]* | Genetic cross | In this study |
| TXJ0230 | *xjEX0007[Pstr-1::nmtn-1+Pmyo-3:: mCherry]* | Microinjection | In this study |
| TXJ0237 | *xjEX0008[Pstr-1::nmtn-1+Pmyo-3:: mCherry]* | Microinjection | In this study |
| TXJ0228 | *xjEX0009[Pstr-1::nmtn-1+Pmyo-3:: mCherry]* | Microinjection | In this study |
| TXJ0235 | *xjEX0010[Pnmtn-1::nmtn-1+Pmyo-3:: mCherry]* | Microinjection | In this study |
| TXJ0486 | *xjEx0019[Pnmtn-1::MNG::NMTN-1+P str-1:: mCherry +Plin-44::GFP]* | Microinjection | In this study |
| TXJ0571 | *xjEx0031[Pstr-2::GFP+Pmyo-3:: mCherry]* | Microinjection | In this study |
| TXJ0570 | *nmtn-1(tm5438) I; xjEx0031[Pstr-2::GFP+Pmyo-3:: mCherry]* | Genetic cross | In this study |
| TXJ0912 | *xjEx0072[Pstr-1::OSM-6::MNG+Pstr-1::mCherry+Plin-44::GFP]* | Microinjection | In this study |
| TXJ1187 | *nmtn-1(tm5438) I; xjEx0072[Pstr-1::OSM-6::MNG+Pstr-1::mCherry+Plin-44::GFP]* | Genetic cross | In this study |
| TXJ1000 | *xjEx0073[Pnmtn-1::NMTN-1::MNG+P str-1::mCherry+Plin-44::GFP]* | Microinjection | In this study |
| TXJ1015 | *xjEx0074[Pstr-1::TAX-4::sfGFP+ Pstr-1::mCherry+Plin-44::GFP]* | Microinjection | In this study |
| TXJ1206 | *nmtn-1(tm5438) I; xjEx0074[Pstr-1::TAX-4::sfGFP+ Pstr-1::mCherry+ Plin-44::GFP]* | Genetic cross | In this study |

| | | | |
|---|---|---|---|
| TXJ1028 | *xjEx0075[Pnmtn-1::GFP+Pstr-1:: mCherry +Plin-44::GFP]* | Microinjection | In this study |
| TXJ1031 | *xjEx0076[Pnmtn-1::GFP+Pstr-2:: mCherry+Plin-44::GFP]* | Microinjection | In this study |
| TXJ1030 | *xjEx0077[Pnmtn-1::GFP+Podr-10:: mCherry+Plin-44::GFP]* | Microinjection | In this study |
| TXJ1029 | *xjEx0078[Pnmtn-1::GFP+Psrb-6:: mCherry+Plin-44::GFP]* | Microinjection | In this study |
| TXJ1229 | *xjEx0096[Pstr-1::DYF-19::mCherry + Pstr-1::MNG::nmtn-1+Plin-44::GFP]* | Microinjection | In this study |
| TXJ1311 | *xjEx0097[Pstr-1::DYF-19::mCherry+ Pstr-1::GFP+Plin-44::GFP]* | Microinjection | In this study |
| TXJ1260 | *nmtn-1(tm5438) I; xjEx0097[Pstr-1::DYF-19::mCherry+ Pstr-1::GFP+Plin-44::GFP]* | Genetic cross | In this study |
| TXJ1271 | *xjEx0082[Pnmtn-1::GFP+Pmyo-3::m Cherry]* | Microinjection | In this study |
| TXJ1366 | *xjEx0098[Pstr-1::KAP-1::MNG+Pstr-1::mCherry+Plin-44::GFP]* | Microinjection | In this study |
| TXJ1380 | *nmtn-1(tm5438) I; xjEx0098[Pstr-1::KAP-1::MNG+Pstr-1::mCherry+Plin-44::GFP]* | Genetic cross | In this study |
| TXJ1399 | *xjEx0099[Pstr-1::OSM-3::MNG+Pstr-1::mCherry+Plin-44::GFP]* | Microinjection | In this study |
| TXJ1432 | *nmtn-1(tm5438) I; xjEx0099[Pstr-1::OSM-3::MNG+Pstr-1::mCherry+Plin-44::GFP]* | Genetic cross | In this study |
| TXJ1428 | *xjEx0100[Psrb-6::OSM-6::MNG+Plin-44::GFP]* | Microinjection | In this study |
| TXJ1431 | *nmtn-1(tm5438) I; xjEx0100[Psrb-6::OSM-6::MNG +Plin-44::GFP]* | Genetic cross | In this study |
| TXJ1429 | *xjEx0101[Pstr-1::TAX-4::sfGFP+ Pstr-1::DYF-19::mCherry+Plin-44::GFP]* | Microinjection | In this study |
| TXJ1430 | *nmtn-1(tm5438) I; xjEx0101[Pstr-1::TAX-4::sfGFP+ Pstr-1::DYF-19::mCherry + Plin-44::GFP]* | Microinjection | In this study |
| TXJ1045 | *xjEx0079[Parl13::MKS-5::mCherry + Pstr-1::mng::nmtn-1+Plin-44::GFP]* | Microinjection | In this study |
| TXJ1033 | *xjEx0080[Parl13::DYF-19::mCherry + Pstr-1::mng::nmtn-1+Plin-44::GFP]* | Microinjection | In this study |
