## Supplementary Table 2 for "WDR47 facilitates ciliogenesis by modulating intraflagellar transport"

| Supplementary Table 2. Key resources table | | |
|---|---|---|
| **Chemicals and Kits** | | |
| Reagent or Resource | Source | Identifier |
| 2,3-Butanedione monoxime | Sigma | Cat#: B0753 |
| Levamisol hydrochloride | Sigma | Cat#: 31742 |
| Dil | Molecular Probes | Cat#: D-282 |
| 2-Nonanone | Aladdin | Cat#: N105585 |
| Diacetyl | Sinopharm | Cat#: 80042427 |
| Octanol | Sangon Biotech | Cat#: A504032-0250 |
| Isopentyl alcohol | Sangon Biotech | Cat#: A610278-0500 |
| QIAprep Spin Miniprep Kit | Qiagen | Cat#: 27106 |
| TIANPrep Rapid Mini Plasmid kit | TIANGEN | Cat#: DP105-03 |
| PrimeSTAR Max DNA Polymerase | Takara | Cat#: R045A |
| Hieff CLoneTM Plus One Step Cloing Kit | Yeasen | Cat#: 10911ES62 |
| TIANquick Midi Purification Kit | TIANGEN | Cat#: DP204-03 |
| **Primers information** | | |
| P*str-1* F | aagcttgagtgagaagaatgtcacg | |
| P*str-1* R | gtcgactagtcaaatgatatgaagtt | |
| P*str-2* F | aagcttatataaatcaatgggatc | |
| P*str-2* R | gtcgacttttatggatcacgagtattc | |
| P*odr-10* F | aagcttttaatttttcataattgactc | |
| P*odr-10* R | gtcgacggagctgtaaggtatcttaatg | |
| P*srb-6* F | tctacttttaaatattatatctttctaattttttgcaacgaa | |
| P*srb-6* R | ttttatttcttctgtagaaatttcaagactgatca | |
| P*nmtn-1* F | gaacccaattgacataatgctctc | |
| P*nmtn-1* R | tttagcatggataatgttattgcg | |
